# MHC-II molecules present RhoC-derived peptides on the surface of tumour cells

**DOI:** 10.1101/2022.05.15.492002

**Authors:** Pavana Thomas, Sweta Srivastava

## Abstract

RhoC is an important regulator of metastasis and tumour progression across various tumour models. Since RhoC has been found to have no major contribution towards normal embryogenic development, it has emerged as a suitable therapeutic target for effective cancer treatment. Recent evidence has shown that Rho-based peptide vaccines have favourable outcomes in prostate cancer patients, by bringing about activation of CD4^+^ T-cells. Antigen presentation on the surface of cells is brought about by the MHC-I/MHC-II complex. This work provides conclusive evidence to show that the seemingly cytosolic protein, RhoC, is in fact, present on the surface of tumour cells. This report goes on to prove that the presentation of RhoC peptides is brought about in association with MHC-II, becoming the first piece of scientific evidence to report this phenomenon.

## INTRODUCTION

It has become increasingly evident that metastasis contributes to poor clinical outcomes (1). The progress from primary tumour to secondary metastatic outgrowths involves cruising of cancer cells through dynamically changing microenvironments (2). Essentially, a cell having the ability to overcome and adapt to the challenges posed by changing microenvironments eventually form metastases (3). Due to these factors, the molecular landscape of primary tumour cells and metastatic cells are completely different, and the therapeutic strategies used to target these two cell types must therefore be unique.

RhoC has been shown to regulate tumour progression in several tumour models (4–7). It has been shown to modulate several hallmarks of cancer including invasion, proliferation, anoikis resistance, survival, tumorigenicity, therapy resistance and metastasis (8). We have recently reported that RhoC regulates therapy resistance in cervical cancer (9). Collectively, data positions RhoC as an important regulator of tumour progression and metastasis. Importantly, it must be noted that RhoC has been shown to be essential for metastasis but dispensable for murine embryogenesis (10). Additionally, mutation analysis reveals that RhoC does not have mutations which are driver, and may only have passenger mutations, suggesting that these genetically variant clones may not evolve due to therapy selection pressure (11). It was thus important to develop targeted therapy against RhoC.

In this regard, peptide-based vaccination as immunotherapy has been explored using RhoC as a target. Reports suggest that HLA-A3 restricted epitope of RhoC is recognized by cytotoxic T-cells, and that RhoC-specific T-cells exhibit cytotoxic activity against cancer cells of varied origins with matched HLA types (12). Further, phase I/II clinical trials on the said peptide suggests that a majority of the patients developed enhanced CD4^+^ T-cell response (13).

Canonical pathways suggest that MHC II (HLA-class II) complex activity is involved in CD4^+^ T-cell activation. Antigen presenting cells are dominant activators of CD4^+^ T cells as they express the MHC-II complex (14). All nucleated cells on the other hand, primarily express the MHC-I complex (15). Thus, it was exciting to understand the mechanism of presentation of RhoC peptides.

The presentation of peptides by antigen presenting cells resulting in activation of CD4^+^ T cells, and consequent targeting of tumour cells via activation of CD8^+^ cytotoxic T cells is an expected paradigm. However, it raises the question of recognition of tumour cells. Do the tumour cells have RhoC on the cell surface, and if so, how are they presented? It is to be noted that RhoC is a cytosolic protein with newly reported nuclear functions, but no cell surface expression of this protein has been reported till date. Thus, in the present study, we examined the new paradigm of presence of RhoC on the surface of tumour cells, and the mode of presentation, which may aid identification by CD4^+^ T-cells and CD8^+^ cytotoxic T-cells.

## MATERIALS AND METHODS

### Cell culturing

The cell lines were cultured in their recommended media (as given in the table below), supplemented with 10% FBS (Foetal Bovine Serum) and penicillin-streptomycin (Invitrogen Catalogue number:15140148) at 37°C and 5% CO_2_. The media used were as follows-

| **Cell line** | **Media used** |
| --- | --- |
| SiHa | DMEM |
| PC-3 | RPMI |
| Huh7 | DMEM-F12 |
| A375 | MEM-α |
| HCT-116 | Mc Coy's |
| Mcf7 | DMEM |

### Immunofluorescent staining (fixed cells)

Cells were washed thrice with 1X PBS to remove media. They were then fixed in 4% PFA at room temperature for 15mins. They were then washed with 1X PBS (x3). 50mM NH_4_Cl was added to the cells and incubated at room temperature for 20mins. The cells were permeabilized using 0.1% Triton-X for 5mins at RT. The cells were washed thrice with 1X PBS and then blocked using a blocking buffer containing 5% BSA and 0.2% fish skin gelatin in PBS for 20mins at RT. Primary antibodies were added (in blocking buffer) and incubated overnight at 4°C. The cells were washed thrice with PBS, and secondary antibodies + Hoechst were added and incubated at RT for 45 min. The cells were washed thrice and mounted using VECTASHIELD anti-fade mounting media. The slides were imaged using the Zeiss 710 confocal microscope. ImageJ software was used for image analysis. The Colocalization plugin was used for analysis of colocalized pixels and the Mander’s scores generated were used to determine the extent of colocalization.

### Immunofluorescent staining (live cells)

Cells were washed thrice with 1X PBS to remove media. They were then fixed in 4% PFA at room temperature for 15mins. They were washed with 1X PBS (x3). 50mM NH_4_Cl was added to the cells and incubated at room temperature for 20mins. The cells were washed thrice with 1X PBS and then blocked using a blocking buffer containing 5% BSA and 0.2% fish skin gelatin in PBS for 20mins at RT. Primary antibodies were added (in blocking buffer) and incubated overnight at 4°C. The cells were washed thrice with PBS, and secondary antibodies + Hoechst were added and incubated at RT for 45 min. The cells were washed thrice and mounted using VECTASHIELD anti-fade mounting media. The slides were imaged using the Zeiss 710 confocal microscope. ImageJ software was used for image analysis. The Colocalization plugin was used for analysis of colocalized pixels and the Mander’s scores generated were used to determine the extent of colocalization.

Details of antibodies used and their dilutions-

| **Antibody** | **Catalogue Number** | **Dilution** |
| --- | --- | --- |
| RhoC | CST, #3430 | 1:100 |
| MHC II | Abcam, #ab55152 | 1:200 |
| Alexa 488, anti-rabbit | Invitrogen, A11034 | 1:500 |
| Alexa 647, anti-mouse | Invitrogen, A21236 | 1:500 |

### Immunostaining for FACS

The cells were detached using 5mM EDTA and spun at 2000rpm for 5mins. For total staining, the cells were fixed in 1%PFA at RT for 5mins. The cells were permeabilized using 0.1% TritonX100 and blocked using 5% FBS and 2% BSA. For surface staining, live unfixed non-permeabilized cells were used. The cells were then incubated in primary antibody at the appropriate dilutions at RT for 2hrs. They were then washed thrice in PBS and incubated with secondary antibody at the appropriate dilutions for 45mins at RT. The stained cells were then washed in PBS and analysed using the FC500.

| **Antibody** | **Catalogue Number** | **Dilution** |
| --- | --- | --- |
| RhoC | CST, #3430 | 1:1000 (for fixed cells)<br>1:200 (for live cells) |
| Alexa 488, anti-rabbit | Invitrogen, A11034 | 1:500 |

### Transfection

Hiperfect (Qiagen Catalogue Number: 301705) reagents were utilized for transfection experiments. All siRNAs were procured from Ambion, Invitrogen. Cells were seeded at 70% confluency on the day prior to commencement of transfection experiments. Transfection complexes comprising of siRNA and Hiperfect transfection reagent (3μl for a 35mm dish) were made in an eppendorf tube in 100μl incomplete media (devoid of FBS). The complexes were allowed to stand at room temperature for 10mins. Media was replaced with 900μl of fresh complete media and the complexes were added onto the cells. The dishes were swirled to allow homogenous distribution of the reagents and incubated at 37°C and 5% CO2. Cells were retrieved 48-72h post transfection.

The siRNAs and the quantities used per ml of the media are indicated in the table below-

| Target gene | Quantity (in picomoles) | Catalogue Number |
| --- | --- | --- |
| RhoC | 30 | 120759, 120897 |
| Scrambled (negative control) | At the same concentration as the target siRNA. | AM4611 |

### RNA Isolation

The media was removed and cells were washed thoroughly with PBS. The PBS was removed completely and 1ml of TRIzol (Invitrogen Catalogue Number: 15596026) was added per 10^7^ cells. Cells in TRIzol can be stored at −30°C for upto 6 months. RNA was then isolated as per the protocol detailed by the manufacturer. The isolated RNA was quantified using Qubit and stored at −80°C for further use.

### cDNA preparation

The RNA isolated was converted to cDNA for further analysis by PCR. The MMLV-Reverse transcriptase kit (Catalogue Number: 28025013) from Invitrogen was utilized for this purpose. 1μg of RNA was converted to cDNA. 0.8μl of random hexamers (Invitrogen Catalogue Number: 48190011) at a working concentration of 300ng/μl was added to 1μg RNA in a 0.2ml PCR vial. The volume was made up to 12.5μl with nuclease-free water. The mixture was incubated at 70°C for 10mins. 7μl of a master mix containing the components listed in the table below was added-

| Component | Volume (in µl) |
| --- | --- |
| 5X First Strand Buffer | 4 |
| 0.1M Dithiothreitol | 2 |
| 10mM dNTPs (Invitrogen Catalogue Number: R0181) | 1 |

The tube was then incubated at 37°C for 2mins. 0.5μl of the MMLV-RT enzyme was added into the tubes and incubated under the conditions detailed in the table below.

| Temperature | Time |
| --- | --- |
| 25°C | 10mins |
| 37°C | 60 mins |
| 70°C | 15mins |

The prepared cDNA was stored at −30°C.

### Real Time quantitative PCR

The TB GreenTM Premix Ex TaqTM II mix (Catalogue Number: RR820A) from TAKARA was used to analyse changes in gene expression. The components and the volumes of reagents added are given in the table below-

| Components | Volumes |
| --- | --- |
| Nuclease-free water | 3µl |
| Forward primer (10µM) | 0.5µl |
| Reverse primer (10µM) | 0.5µl |
| Template cDNA | 1µl |
| SYBR Green master mix | 5µl |
| **Total** | 10µl |

The samples were run on a 7500 Fast Real Time PCR machine from Applied Biosystems using the ΔΔCt method. The cycling conditions used are given below-

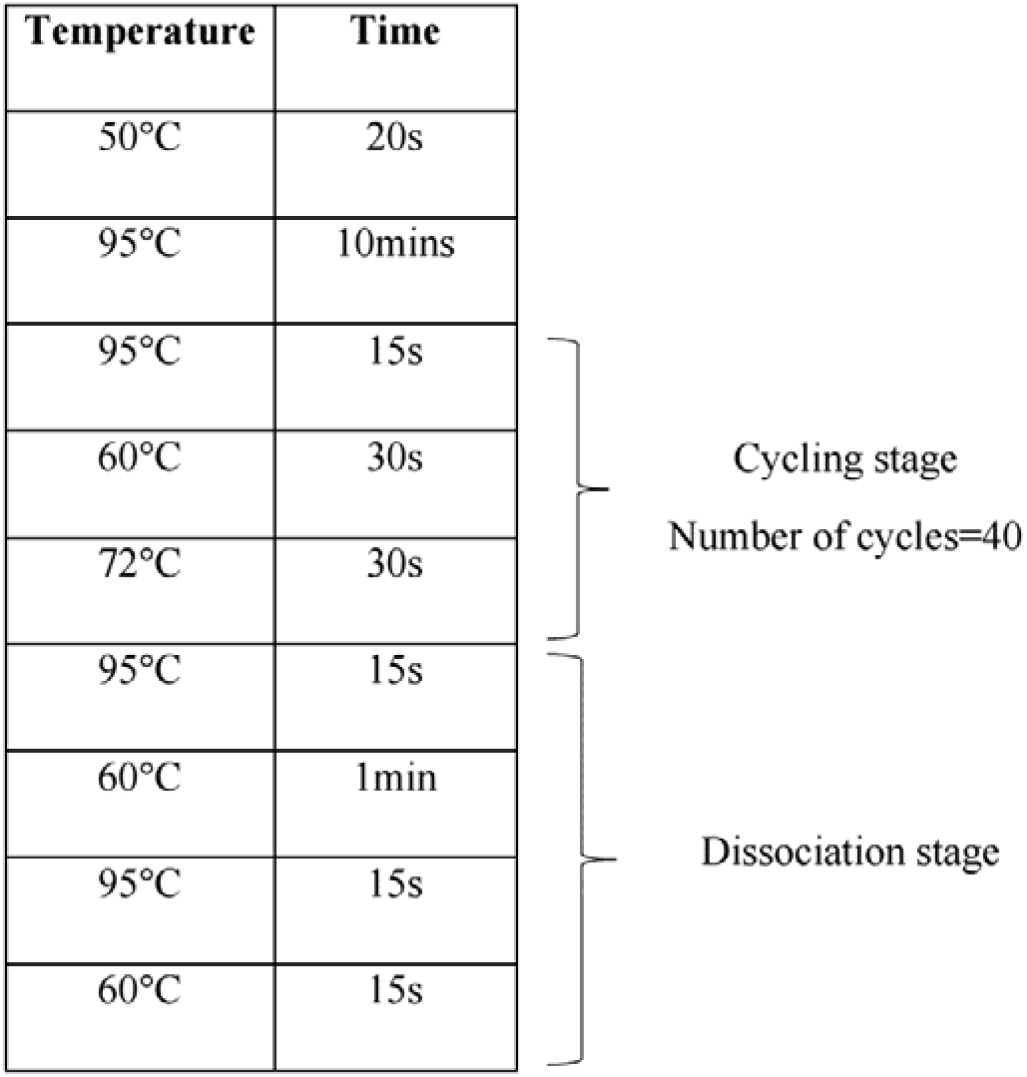

GAPDH was used as the internal control. Samples were normalized to GAPDH and the dCt values thus obtained were used to compute fold change in gene expression.

The sequences of the primers used are detailed in the table below-

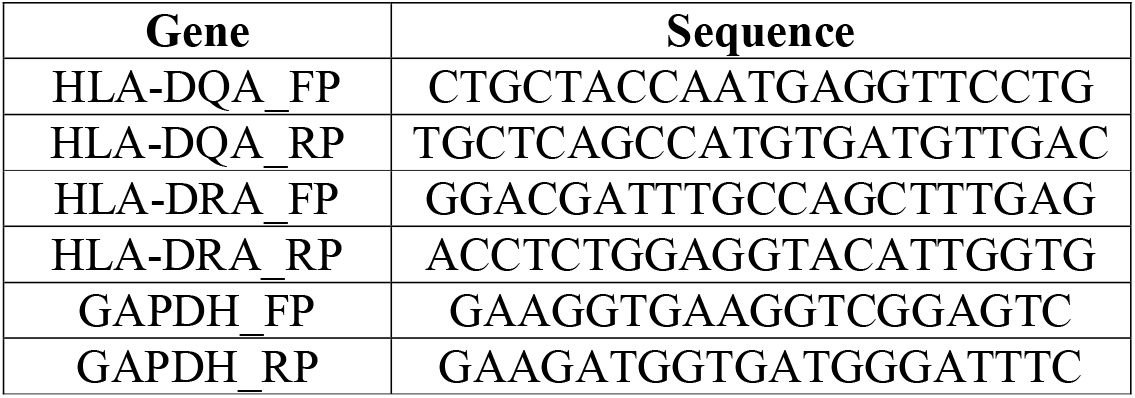

### Immunoprecipitation

Cells were washed with PBS and cross-linked using Lomant’s reagent. The cells were washed gently with cold PBS and then incubated with lysis buffer (50mM Tris HCL (pH7.4), 150 mM NaCl, 2mM EDTA (pH8.0), 1% NP-40, 100mM NaF, 200mM Na3VO4, 10X MPI, 200mM PMSF). The lysate was collected and spun at 13,000 rpm for 15mins at 4°C. The supernatant thus obtained was pre-cleared using equilibrated Protein G dynabeads. The dynabeads were incubated with 2μg of the desired antibody (MHC II (a), Abcam, #ab55152; MHC II (b), Santa Cruz, #32247) at room temperature for 10mins and then added to the precleared lysate. The proteins were allowed to bind overnight under gentle agitation at 4°C. The beads were washed and the protein complexes attached to these beads were subject to dot blot analysis and MS-based identification.

### Dot blot analysis

3μl of the pulled-down complexes were spotted onto a strip of nitrocellulose membrane and allowed to dry. Once dry, the blot was washed thrice in 1X TBST for 10mins each. The blot was then incubated with the required primary antibody at room temperature for 2hrs. The blot was washed thrice with 1X TBST and incubated with secondary antibody at room temperature for 1hr. After three washes with 1X TBST, the blot was developed using ECL reagent and visualized under the Bio-Rad ChemiDoc XRS^+^ system.

### MS/MS identification of proteins

The pulled-down peptides were subjected to MS/MS using the Bruker Daltoincs ESI Q-TOF system with the Proxeon EASY-nLC. The data generated was analysed using the MASCOT database to identify the precipitated proteins.

## RESULTS

### Presence of RhoC on the surface of tumour cells

Recognition of tumour cells by the immune system is the first step towards targeting the former. Since phase I/II trials of RV001, a vaccine comprising of a peptide of the RhoC protein, shows activation of CD4^+^ T-cells (13), it was hypothesized that RhoC may be present on the surface of tumour cells. Thus, to investigate the presence of RhoC on the surface of tumour cells, immunofluorescent-based expression analysis was performed on cells under both permeabilized (PM) and non-permeabilized (NPM) conditions. The expression was analysed using flow cytometry and confocal microscopy to assess the quantitative changes and localization of the protein respectively. We have assessed the presence of RhoC protein on the surface of tumour cells from various tumour models including cervical, prostate, skin, liver, colon and breast cancer. As shown in **Figures 1A(i-vi)**, FACS analysis revealed the presence of RhoC in a sub-population of cells in all the tumour models studied. The percentage of positivity on the surface of these cells ranged from 11-45%. Representative FACS plots for the respective non-permeabilized (NPM) secondary controls are shown in **Figure S1A**. Concurrently, FACS analysis was also performed on these cell lines under permeabilized (PM) conditions **(Figure S1B)**. The percentage of total RhoC positivity, as found in cells under the PM conditions, was found to range from 35-90%. Significantly, it was found that all the cell lines studied displayed greater total RhoC positivity as opposed to surface positivity of RhoC **(Figures 1B(i-vi)**). The presence of RhoC on the surface was further confirmed by confocal microscopy. Confocal imaging revealed positive RhoC expression on the surface of all the cell lines studied **(Figure 1C)**. Additionally, cells under PM conditions were also found to be RhoC positive, confirming the FACS data **(Figure S2)**. Interestingly, it was observed that the level of RhoC on the surface (NPM) reduced upon siRNA knockdown of RhoC **(Figures 1D, 1E, S2B)**. The knockdown of RhoC by siRNA was confirmed by staining cells for total RhoC (PM) **(Figure S2C)**.

**Figure 1:**
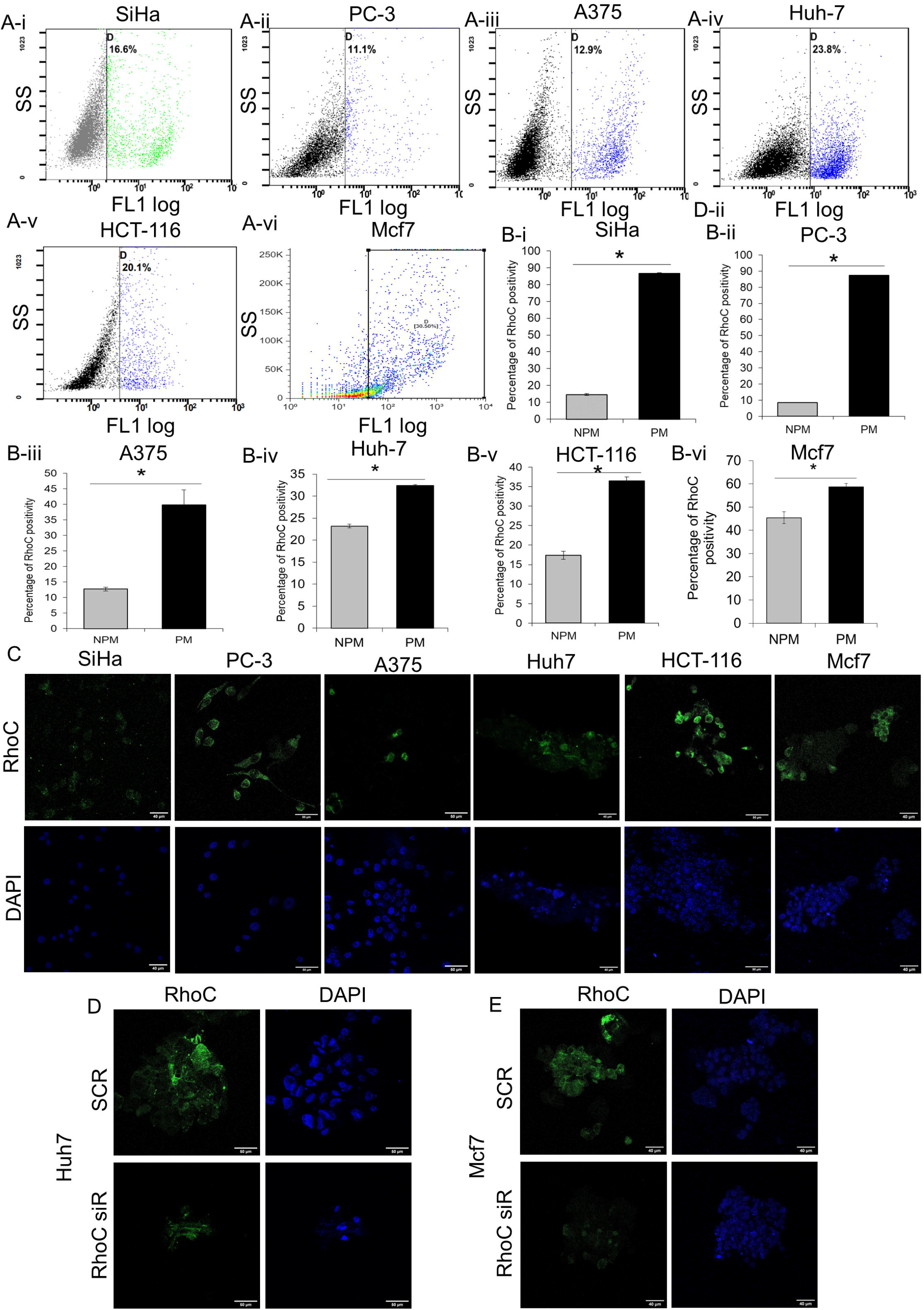
Cytosolic RhoC is detected on the surface of numerous tumour cell lines. **A (i-vi)-** Representative FACS plots showing RhoC positivity in live, non-permeabilized tumour cell lines. SiHa, PC-3, A375, Huh-7, HCT-116 and Mcf7 cells were observed to have a percentage of the population that had surface positivity of RhoC (n=3). **B (i-vi)** -Graphical representation of the percentage of RhoC positivity in non-permeabilized (NPM) and permeabilized (PM) SiHa, PC-3, A375, Huh-7, HCT-116 and Mcf7 cells (n=3). **C-** Confocal microscopy of non-permeabilized cells showing positive staining of RhoC on the surface of SiHa, PC-3, A375, Huh-7, HCT-116 and Mcf7 (n=3). **D-** Confocal microscopy images showing reduction in surface levels of RhoC upon knockdown of RhoC by siRNA transfection in Huh-7 cells (n=3). **E-** Confocal microscopy images showing reduction in surface levels of RhoC upon knockdown of RhoC by siRNA transfection in Mcf7 cells (n=3).

### Expression of MHC II in tumour cells

MHC II is predominantly expressed on antigen presenting cells. However, recent scientific advances and reports suggest that the MHC II complex is over-expressed in tumour cells (16). Further, evidence suggests the activation of the CD4^+^ pool of T cells upon vaccination with the RhoC peptide (13). Investigations were thus made to assess the expression of MHC II in the cell lines studied above for the presence of surface RhoC. qPCR-based analysis showed the presence of transcripts of the alpha chains of DQ and DR-two major alleles of MHC II in all the cell lines assayed **(Figure 2A)**. To confirm the presence of MHC II protein within these cells, immunofluorescent staining was performed. Confocal imaging revealed the presence of MHC II within these tumour cell lines, confirming the qPCR data **(Figure 2B)**.

**Figure 2:**
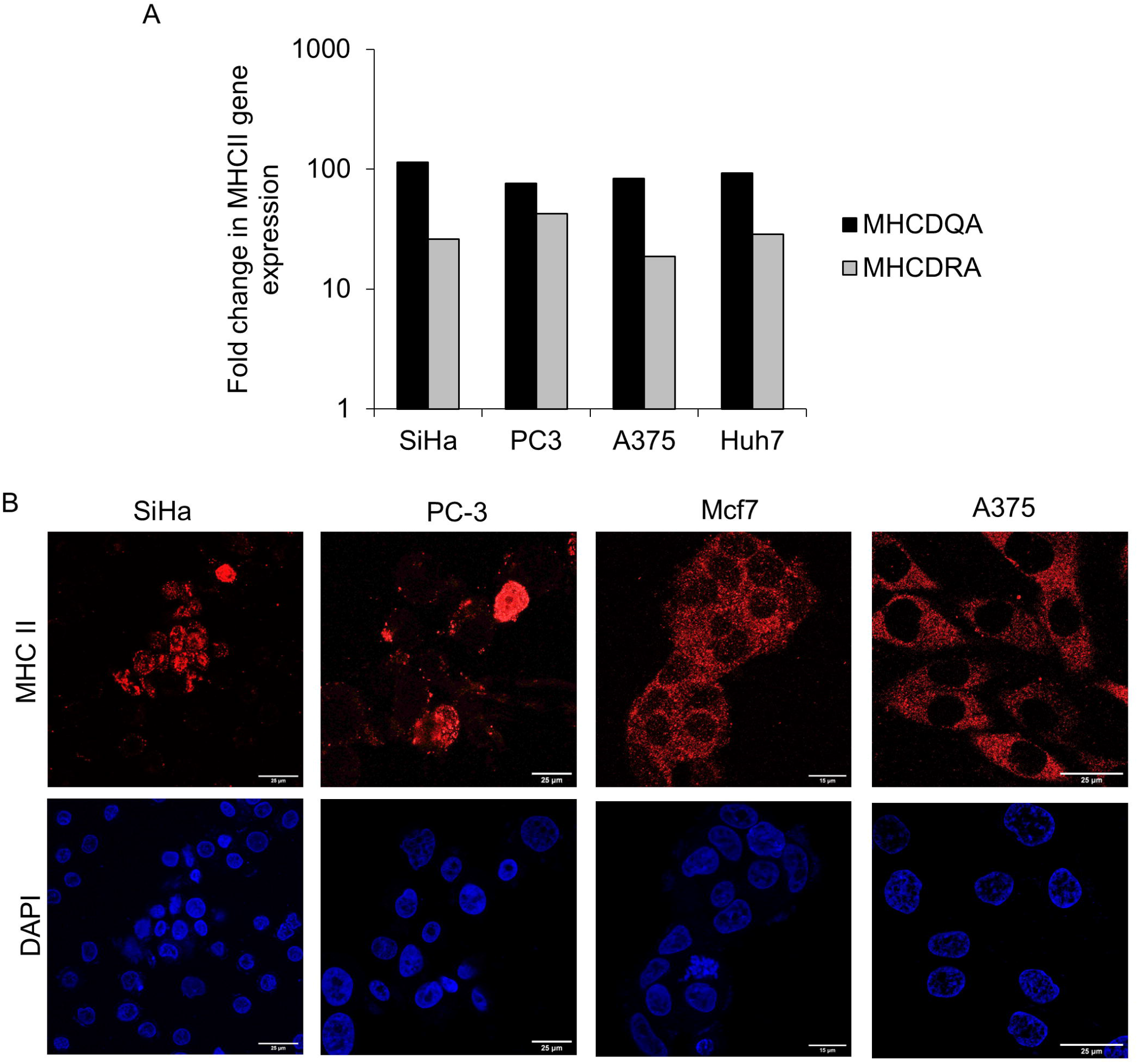
Presence of MHC II in tumour cell lines. **A-** Transcript analysis by qPCR shows the presence of mRNA of MHC DQA and MHC DRA genes (MHC II alleles) in tumour cell lines SiHa, PC-3, A375 and Huh7. **B-** Confocal microscopy shows the presence of MHC II in SiHa, PC-3, Mcf7 and A375 cells (n=3).

### MHC II binds to RhoC peptides and presents it on the cell surface

Studies were done to understand whether MHC-II was involved in RhoC presentation at the cell surface. Confocal microscopy-based colocalization analysis was performed on various tumour cell lines to determine colocalization between RhoC and MHC II. Interestingly, RhoC and MHC II were found to colocalize as observed by the presence of white pixels in the merged image. The Mander’s coefficients calculated by Image J analysis also showed significant colocalization between these two proteins under both NPM and PM conditions **(Figures 3A and 3B)**.

**Figure 3:**
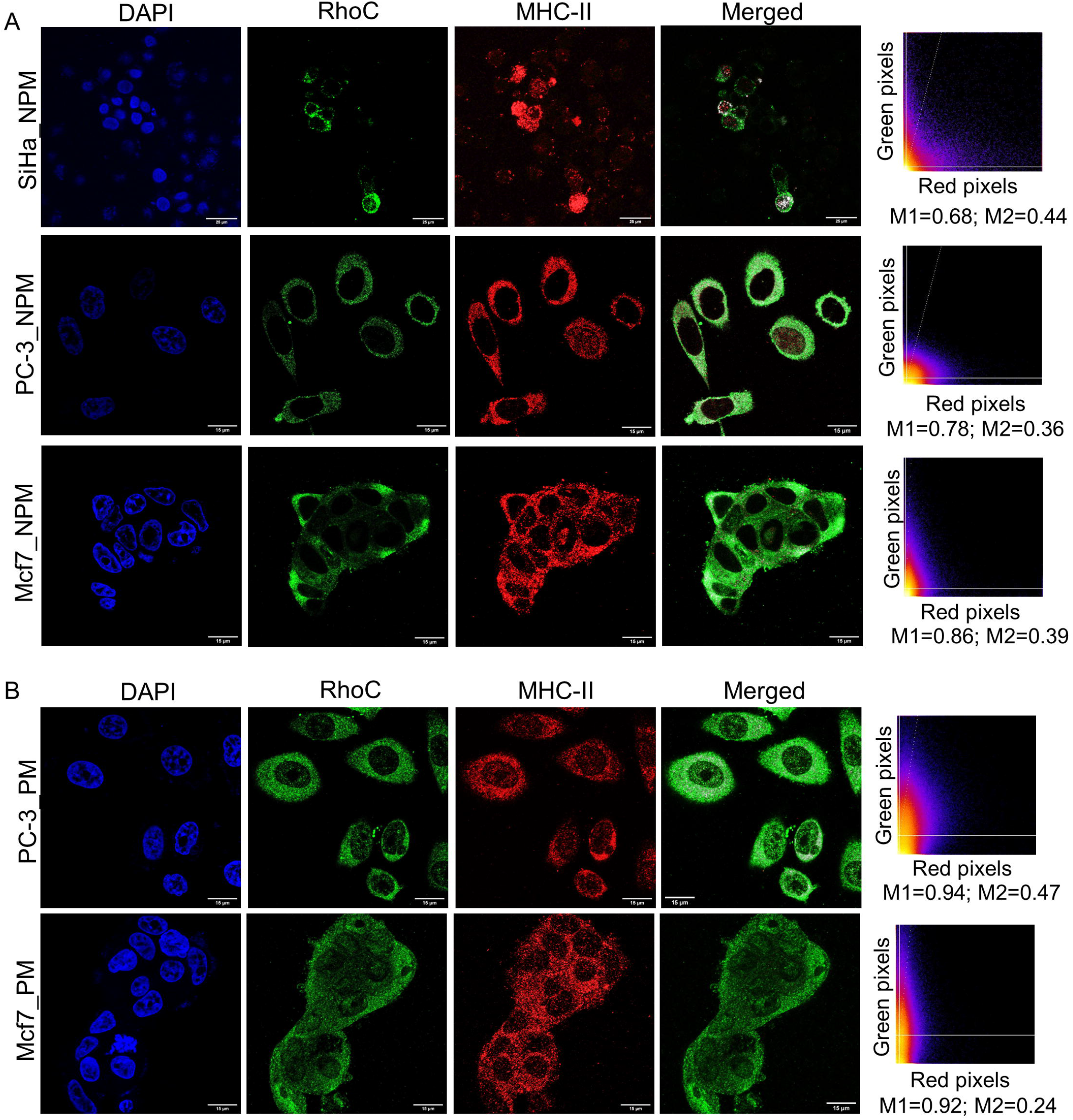
RhoC and MHCII colocalize in tumour cells. **A-** Representative confocal images show significant colocalization of RhoC and MHC II in non-permeabilized (NPM) SiHa, PC-3 and Mcf7 cells. Mander’s coefficients (M1 and M2) signifying the extent of colocalization of red (M1) and green (M2) pixels are also indicated (n=3). **B-** Representative confocal images show colocalization of RhoC and MHC II in permeabilized PC-3 and Mcf7 cells. Mander’s coefficients (M1 and M2) are also indicated (n=3).

### Interaction between MHC II and RhoC

To further confirm that RhoC is presented on cell surface of tumour cells by MHC II, MHC II antibody was used to immunoprecipitate complexes of MHC II and peptides that associated with MHC II. The pulled-down complexes were then subjected to dot blot analysis. Since the RhoC protein was expected to be a small fragment length, conventional PAGE was not used, instead a dot blot was performed to assess binding between the proteins. RhoC was detected in MHC II pull-down complexes as shown in **Figure 4A (i)**, confirming their interaction. Further, MHC II was also detected in the MHC II pull-down, signifying successful immunoprecipitation **Figure 4A (ii)**. Additionally, LC MS/MS analysis was performed to reconfirm this finding and identify the specific peptides of RhoC that associated with MHC II. Analysis of the generated spectral dataset against the human protein database revealed the presence of two peptides of RhoC that interacted with MHC II. The peptides found in close association with MHC II were LVIVGDGACGK and HFCPNVPIILVGNKK. The spectral images are shown in **Figures 4B-i and 4B-ii**. This data thus confirms that the MHC II complex is indeed involved in presentation of peptides of the RhoC protein on the surface of tumour cells.

**Figure 4:**
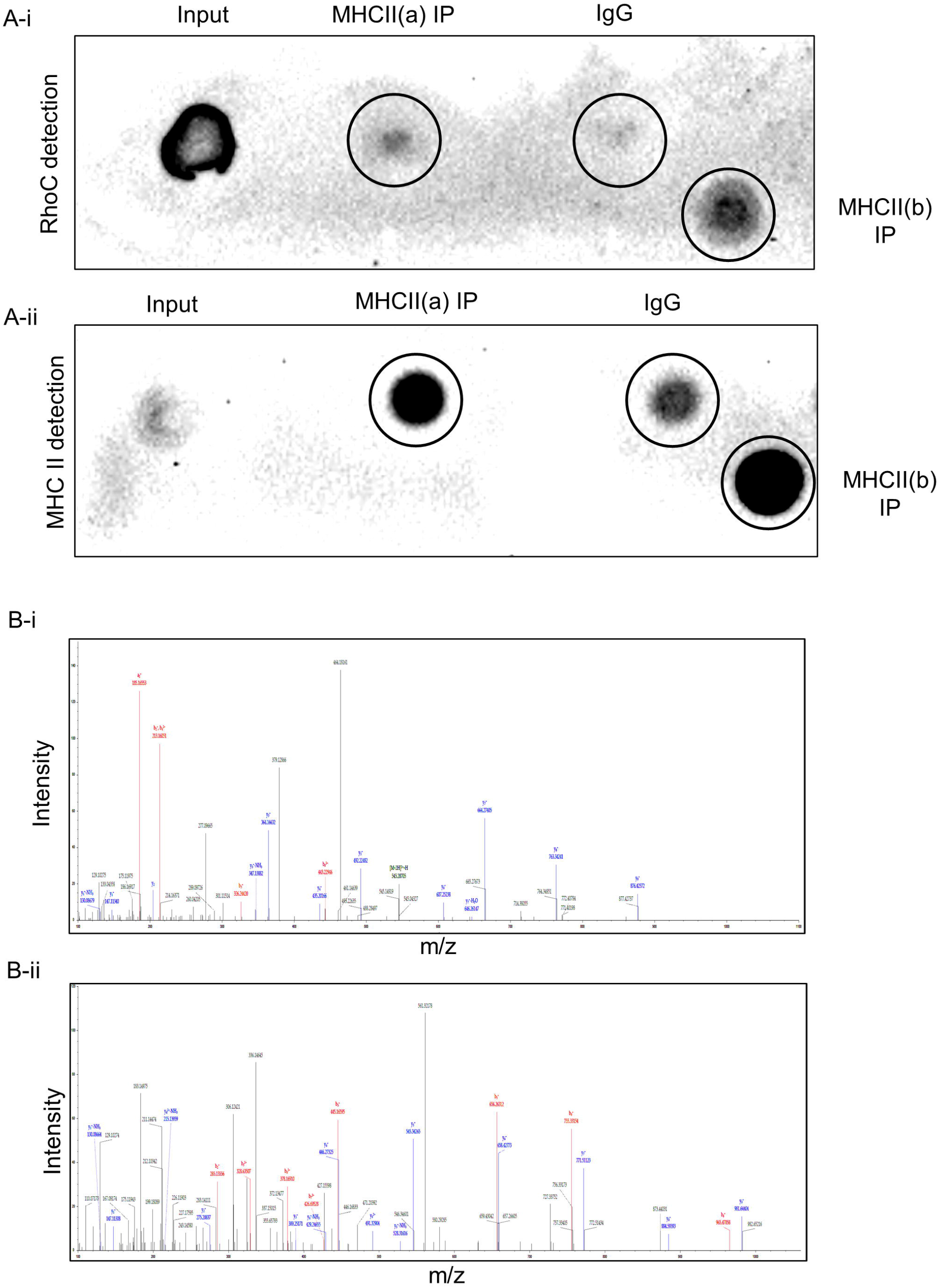
RhoC and MHCII proteins interact with each other in tumour cells. **A (i)-** Dot blot analysis of MHC II immunoprecipitated proteins indicate strong interaction between RhoC and MHC II, as evident by the presence of RhoC in MHC II pull-down complexes. **A (ii)**-Dot blot analysis also confirms successful pull-down indicated by the presence of MHC II in MHC II immunoprecipitated samples. **B (i-ii)**- Spectral peak images of peptides of RhoC identified in MHC II pull-down complexes as confirmed by mass spectrometry analysis (n=2).

## DISCUSSION

Immuno-surveillance and response have been the one among the most important features of tumour life. Immune evasion and escape results in progression of the tumour (17). With the advent of the gene-based therapy model, immunotherapy for treatment of cancers has gained traction (18). The success of immunotherapy has prompted interest in areas of research that explore various immune-related processes for better clinical outcomes. Research efforts have since been directed towards identification of novel biomarkers for immunotherapy and understanding molecular heterogeneity. It is important to note that once a tumour has metastasized, successful therapy becomes a challenge. Therefore, efforts should be directed towards developing targeted therapeutic strategies for metastasis.

Research findings over the past few years have promoted confidence in developing RhoC as a molecular target with therapeutic implications. RhoC, implicated in several phases of tumour progression (19), and tumour phenotypes including invasion (20), angiogenesis (21, 22), anoikis resistance (4), EMT (23–25) and therapy resistance (9) has a significant role in metastasis, and is also a regulator of cancer stem cells (8). The development of a long peptide vaccine targeting RhoC has shown promising outcomes. Phase I/II clinical trials show that patients developed CD4^+^ T-cell response (13). Although RhoC is a cytoplasmic protein, the intriguing result of Phase I/II clinical trials suggests immune recognition of RhoC on the cell surface of tumour cells.

Our data presented here shows that RhoC is indeed present on the cell surface of a variety of tumour cell lines, including liver, colon, cervical, skin, prostate and breast cancer cell lines. This is the first report which suggests that RhoC is present on the cell surface. Classically, RhoC is cytoplasmic protein and has been recently shown to be expressed in the nucleus (9), but no report has been published illustrating its presence on the cell surface.

Although tumour cells are widely known to express MHC-I, there are also reports that show that MHC-II is expressed by a subset of tumours (16). The presentation of peptides on cell surface for immune response is dependent on MHC-I and MHC-II complex. Although tumour cells are widely known to express MHC-I, there are subsets of tumours which overexpress MHC-II. Balasubramaniam et. al, show that MHC-II genes were significantly up-regulated in HPV-associated cervical carcinogenesis (26). Contrary to this, a study on lung cancer reports significant down-regulation of MHC-II in tumour cells (27). Similarly, MHC-II was shown to be aberrantly expressed in triple negative breast cancer tissues using RNA-seq (28). In the present study we observed similar expression of MHC-II in cancer cell lines including SiHa, PC-3, A375 and Mcf7.

Expression of MHC-II protein in triple negative breast cancers is indicative of good prognosis and lymphocyte infiltration. This may be due to antigenic presentation of tumour antigens on the cell surface resulting in immune activation (29). Conforming to a similar phenomenon, we report that RhoC is presented on the surface of tumour cells and associates with MHC-II proteins. This was further confirmed by immunoprecipitation of MHC-II which co-immunoprecipitates RhoC peptides, suggesting that MHC-II protein is responsible for RhoC peptide presentation on the cell surface to enable immune response and CD4^+^ T cell activity.

## ACKNOWLEDGEMENTS

We would like to thank RhoVac ApS, Denmark for providing the funding for this work.

