## Supplementary Figures for "MHC-II molecules present RhoC-derived peptides on the surface of tumour cells"

### Slide 1
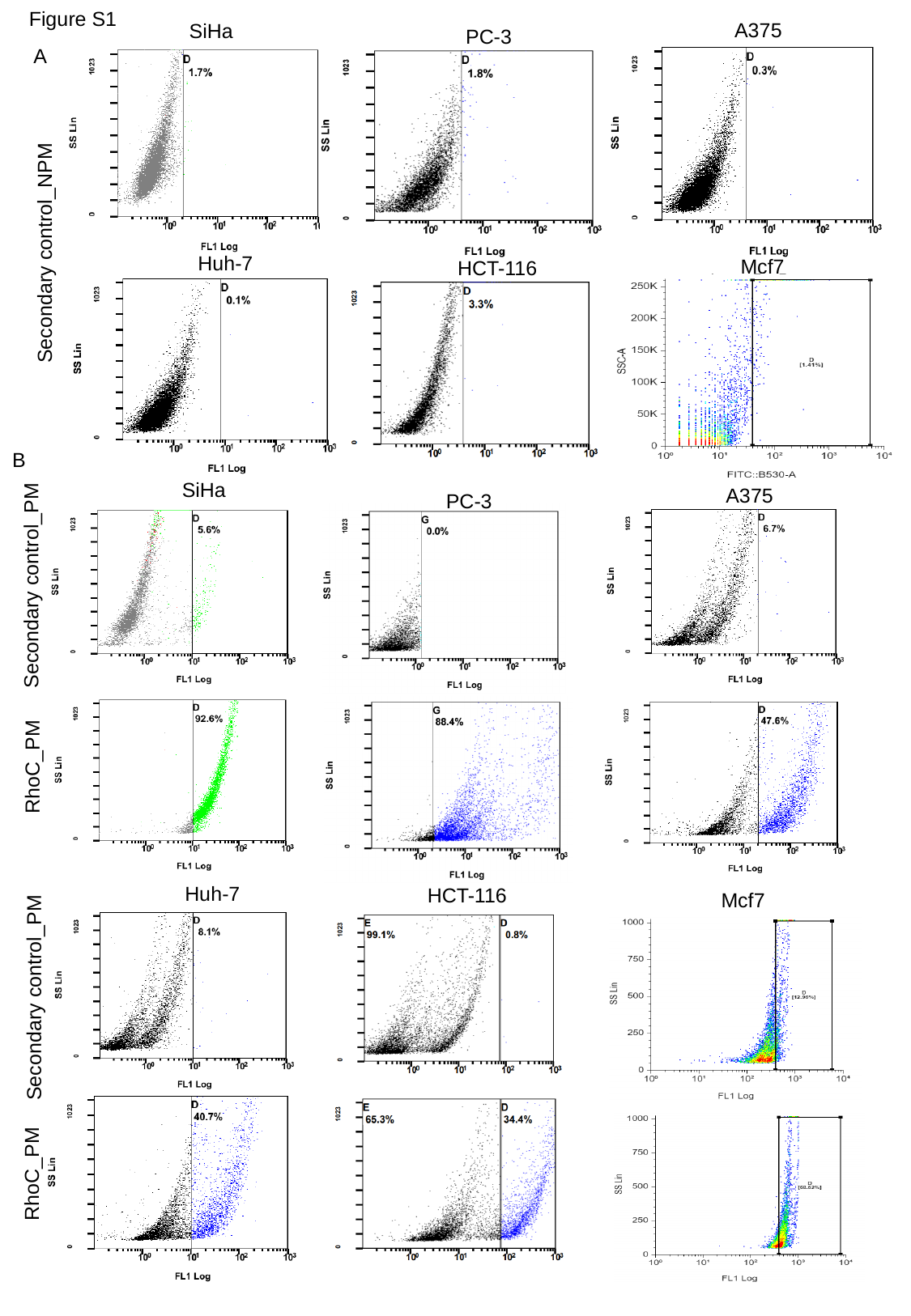

Figure S1
A375
SiHa
PC-3
A
Secondary control_NPM
Huh-7
Mcf7
HCT-116
B
SiHa
A375
PC-3
Secondary control_PM
RhoC_PM
Huh-7
HCT-116
Mcf7
Secondary control_PM
RhoC_PM

### Slide 2
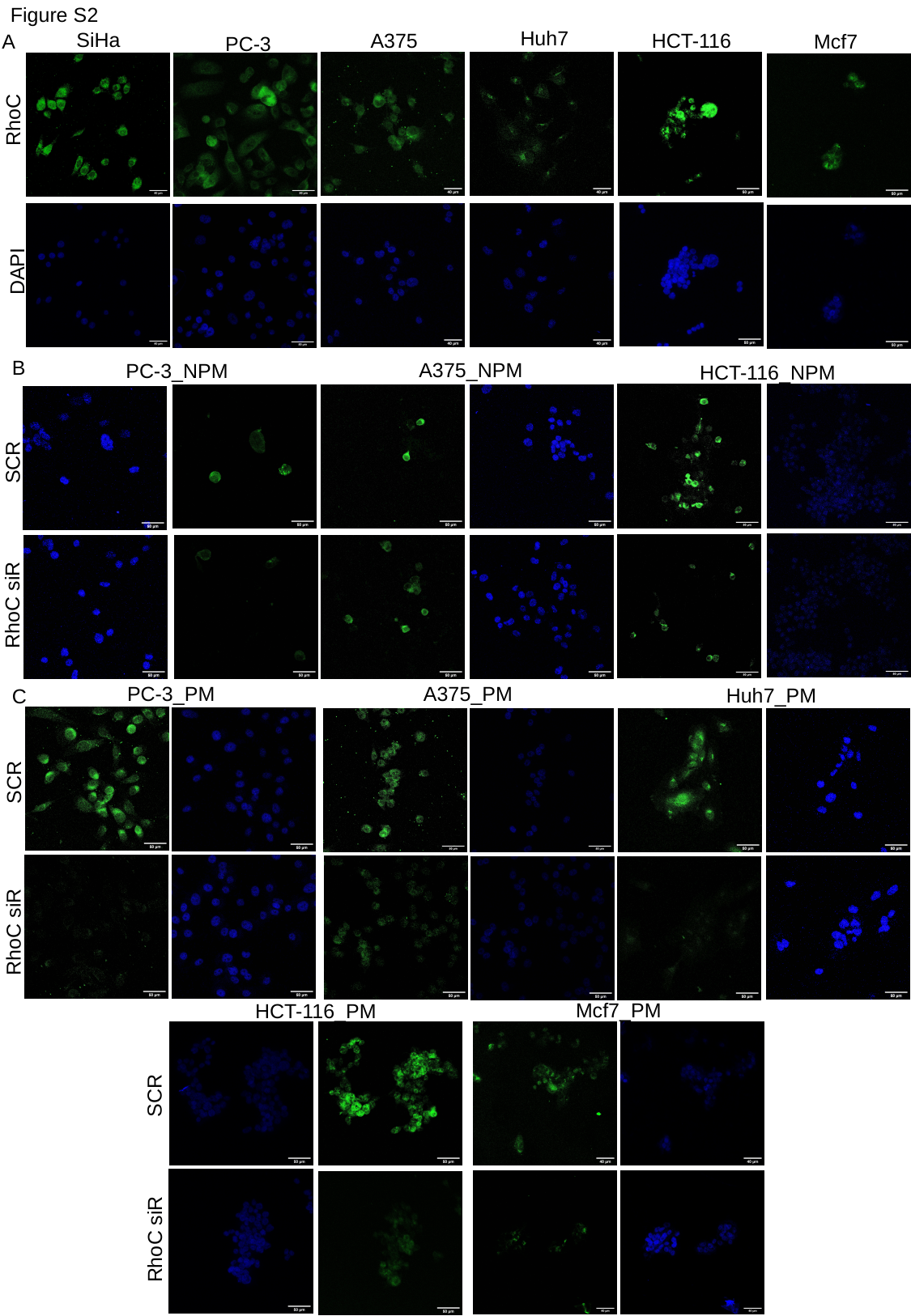

Figure S2
Huh7
SiHa
RhoC
DAPI
HCT-116
A375
A
Mcf7
PC-3
B
A375_NPM
PC-3_NPM
SCR
RhoC siR
HCT-116_NPM
PC-3_PM
SCR
RhoC siR
A375_PM
Huh7_PM
C
Mcf7_PM
HCT-116_PM
SCR
RhoC siR
