## Supplementary figure legends for "MHC-II molecules present RhoC-derived peptides on the surface of tumour cells"

**Figure S1: A-** Representative FACS plots for secondary controls in non-permeabilized (NPM) SiHa, PC-3, A375, Huh-7, HCT-116 and Mcf7 cells (n=3). **B-** Representative FACS plots for RhoC in permeabilized (PM) cells along with respective secondary controls (n=3).

**Figure S2: A-** Representative confocal images for RhoC staining in PM cells (n=3). **B-** Representative confocal images of surface RhoC staining in NPM cells showing reduction in surface RhoC levels upon RhoC knockdown in PC-3, A375 and HCT-116 cells (n=3). **C-** Representative confocal images of surface RhoC staining in NPM cells showing reduction in RhoC levels upon RhoC knockdown in PC-3, A375, HCT-116, Huh7 and Mcf7 cells (n=3).
